# Multigenerational inheritance of parasitic stress memory in *Drosophila melanogaster*

**DOI:** 10.1101/2023.10.11.561819

**Authors:** Shagufta Khan, Ravina Saini, Runa Hamid, Rakesh K Mishra

## Abstract

Organisms sense harmful environmental conditions and employ strategies to safeguard themselves. Moreover, they can communicate this experience to the next generation or beyond via non-DNA sequence-based mechanisms, referred to as intergenerational or transgenerational epigenetic inheritance, respectively. Using a specialist larval parasitoid, *Leptopilina boulardi*, and its host, *Drosophila melanogaster*, we show that the parental experience of parasitic stress results in an increased survivability of the immediate offspring of the host. Furthermore, we observe that the increased survivability in response to the parasitic stress is transmitted transgenerationally where the grandparents have been exposed to the parasitoid but not the parents. The increased survivability is primarily inherited through male parents, and at least one of the forms of the memory is better immune priming at larval stage. Our study suggests that the stress exposure during the pre-adult stage of the host has lifetime benefits for its progeny to deal with the future parasitic attack.

## INTRODUCTION

When faced with detrimental environmental conditions, organisms use strategies to ensure their offsprings’ survival. Since such encounters frequently happen over short evolutionary timescales, the sole role of naturally selected changes in the DNA sequence in providing adaptive plasticity to cope with the challenging environment is difficult to envisage. In such scenarios, however, non-DNA sequence-based multigenerational epigenetic inheritance (MEI) mechanisms, both intergenerational and transgenerational, prove beneficial as they offer a faster and, more importantly, reversible way of imparting adaptive plasticity^1–3^.

Examples of MEI in response to detrimental abiotic factors are plenty but biotic factors, which are equally prevalent, are only recently brought to light^4^. They include non-mutualistic interactions between species, such as parasitism. In parasitism, one organism, referred to as the parasite or pathogen, causes harm to the other organism, the host, by either living on or inside it^5^. As a result of such interactions, adaptive multigenerational epigenetic effects encompassing both behavioral and physiological defences have been reported in a wide range of taxa, such as bees^6,7^, pipefish^8–10^, ragworm^11^, honeycomb moths^12^, *Drosophila*^13–15^, beetles^16–20^, brine shrimps^21^, *C. elegans*^22,23^, and mice^24^.

*Drosophila* genus is host to a plethora of parasites in the natural environment, such as viruses, bacteria, fungi, and even insects called parasitoids^25^. Female parasitoid wasps of the *Leptopilina* genus infect the larval stages of *Drosophila.* They oviposit their eggs into the larval hemocoel along with immuno-suppressive factors, such as venom proteins or virus-like particles^26–28^. In case of a successful infection, the developing wasp consumes the host entirely, develops in the host system, and eventually emerges as an adult from the host pupal case. Occasionally, the *Drosophila* larva mounts a successful immune response, kills the developing wasp, and survives to adulthood. Such flies are referred to as escapee flies^28,29^.

Likewise, *Drosophila* species exhibit numerous other physiological and behavioural defences to safeguard themselves against infection by adult wasps, both at pre-adult and adult stages. For instance, when *Drosophila* adults sense the presence of wasps, they either prefer laying eggs in ethanol- or alkaloid-containing food to medicate their offspring against wasp infection at larval stage^13,15,30–33^ or suspend oviposition^34^. Furthermore, oviposition suspension behavior is communicated to naïve individuals in an intra- or inter-specific manner to confer protection against infection^34–36^. They also increase the production of recombinant over non-recombinant offspring^37^, which may impart fitness to the progenies^38^. Additionally, the host can prime the immune system of their offspring upon cohabitation with adult wasps^14^. At pre-adult stages, however, the larvae exhibit rolling behavior to avoid attack by the wasps^39,40^.

One positive outcome arising from the interaction between hosts and their parasitoids is the phenomenon known as immune priming or immune memory. This phenomenon entails an improved immune response upon re-encountering a pathogen or parasitoid^41,42^. In the context of *Drosophila’s* association with its parasitoid, *Leptopilina boulardi*, immune priming denotes the *Drosophila* host’s capacity to mount a more robust and effective immune response when facing the same or a similar parasitoid following an initial exposure. It follows a three-step progression: initial exposure, the formation of immune memory, and the reinforcement of the response during subsequent encounters^14,43^. In the context of invertebrates, there exists empirical evidence substantiating their ability to transmit this specific form of memory to future generations. This transfer involves the inheritance of traits from both male and female lineages, encompassing occurrences within a single generation (intergenerational) as well as extending across multiple generations (transgenerational)^4,42^. This memory could involve the activation of specific immune-related genes, the production of antimicrobial peptides, or other mechanisms that assist the host in recognizing and responding more effectively to *Leptopilina boulardi* in future encounters^44–46^.

In the present study, we have employed the non-mutualistic association between a specialist parasitoid wasp, *Leptopilina boulardi*, and its host, *Drosophila melanogaster* (or fruit flies), to investigate if repeated infection by the parasitoid wasp for multiple successive generations results in immune priming of the host’s offspring. Interestingly, we observe that the offspring from experienced parents show better survival chances for the wasp attack compared to the progenies of naïve parents. Next, we investigated the potential for both parents to independently pass down acquired immune memory to their offspring and observed that the inheritance of enhanced survivability is more pronounced through the paternal lineage; however, the maternal lineage can transmit the memory for only one generation. Moreover, we show that the memory of enhanced survivability is inherited transgenerationally via the male germ line. Lastly, we explored the potential transmission of transgenerational resistance through an immunological response and found that the larvae born to experienced parents displayed heightened lamellocyte levels when exposed to simulated parasitoid challenges compared to naïve progenies. Overall, our observations indicate a positive correlation between enhanced resistance and an augmented immune response in progeny that exhibited both inter- and trans-generational patterns of inheritance.

## MATERIALS AND METHODS

### Fly strain and culture

The wild-type strain of *Drosophila melanogaster*, or fruit fly, named *Canton-S* (*CS*), was used in the current study. Flies were cultured in bottles at a constant average density of 100-150 flies on a standard medium containing corn flour, sugar, yeast, malt, agar, and preservatives. Flies were maintained throughout at 25°C with a 12-hour light-dark cycle.

### Wasp strain and culture

The Lb17 strain of the parasitoid wasp, *Leptopilina boulardi*, used in the study was kindly provided by Shubha Govind (Biology Department, The City College of the City University of New York). Wasps were cultured on the *CS* strain of *D. melanogaster*, as previously described^47^. Briefly, 2-4 day old flies were allowed to lay embryos for 48 hours at 25°C in standard medium vials. Subsequently, flies were removed, and 6 to 8 young pre-mated female and male wasps were added to the vials to infect the 0-48 hour hosts after egg lay (AEL) (second instar fly larvae). Flies that survived the wasp infection (escapee flies) were removed from the vials immediately upon emergence, and vials were kept for further development of wasps. After 20-22 days, freshly eclosed wasps were taken and used for parasitic stress experiments.

### Multigenerational parasitic stress

F_0_ *CS* flies were mated to collect 0-24 hour F_1_ embryos in the food vials. At 24-48 hours AEL, the larvae were exposed to parasitic stress by infecting with 6-8 Lb17 wasps for 24 hours at 25°C. After infection, the wasps were removed, and the infected larvae were allowed to grow until escapee flies and wasps emerged. The F_1_ escapee flies were collected and mated to obtain the 0-24 hour F_2_ embryos. F_2_ progenies were infected at the second instar larval stage to get F_3_ escapee flies and wasps. An identical method of embryo collection and parasitic stress was performed for ten generations based on the scheme presented in Figures 1A and 1B, giving rise to the experienced treatment group.

**Figure 1.**
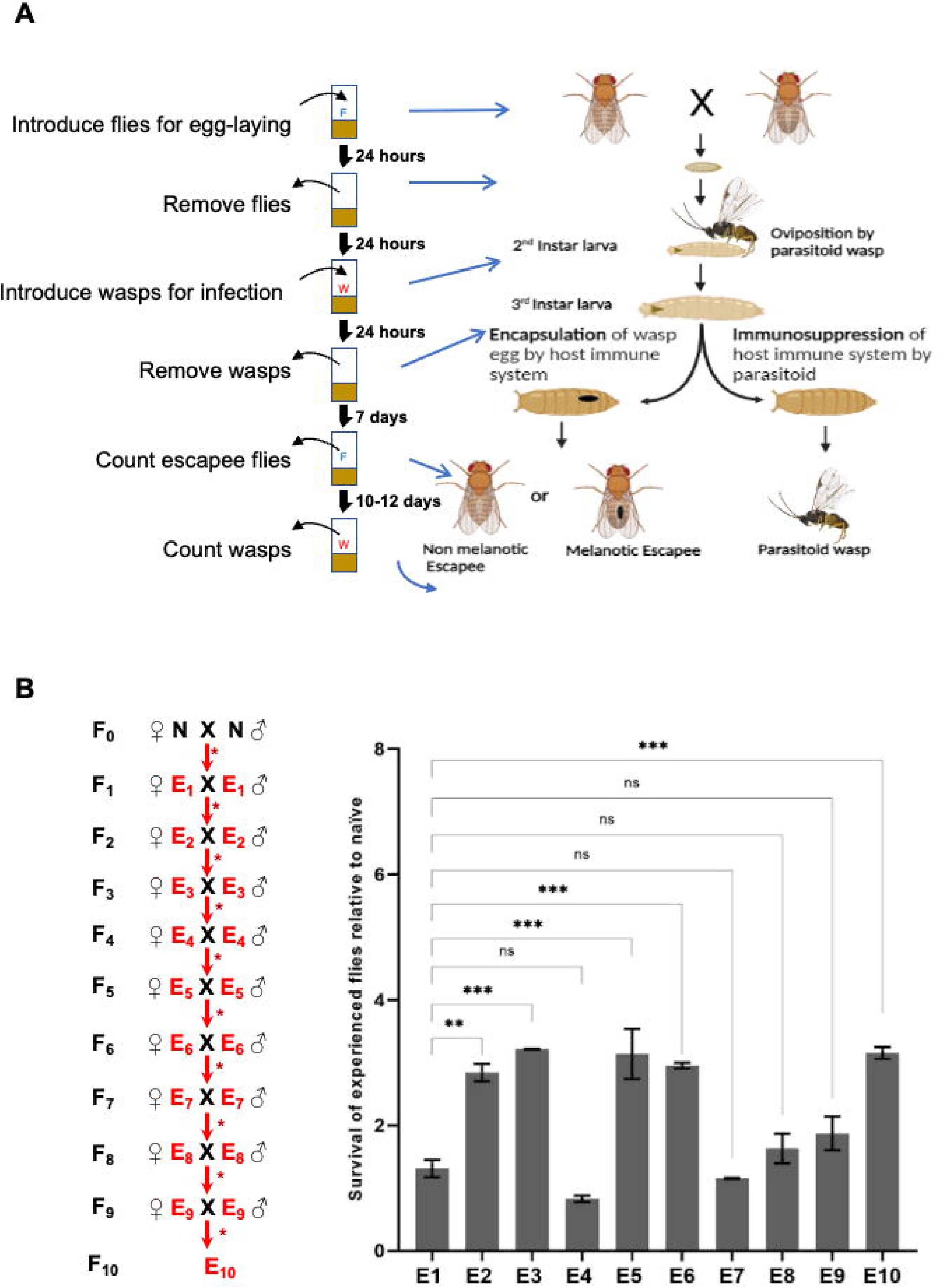
Multigenerational parasitic stress. **(A)** The parasitoid wasp (*Leptopilina boulardi*) lays eggs in the second instar larva of *Drosophila melanogaster*, along with immune-suppressive factors. The first reaction of the host to the wasp egg is to activate cellular immunity and encapsulate it with special immune cells called lamellocytes. Successful hosts emerge as adults (10 days AEL) either with a black melanotic capsule or without a melanotic capsule. If the host fails to encapsulate the parasitoid egg and the parasitoid successfully suppresses the host’s immune system, the parasitoid emerges after 18-20 days of infection. Adult flies that emerge from this host-parasitoid interaction are called experienced escapees (E). W, wasps; F, flies. **(B)** The mating scheme for multigenerational parasitic stress is shown on the left side of the graph. First-time exposed male and female escapees (E_1_) are mated, and their progeny are collected to expose them to obtain second-generation escapees (E_2_). The same scheme is used for up to 10 generations. The text and arrow with an asterisk in red indicate wasp treatment. N, naïve; E, experienced escapee. The survival rate at every generation is calculated and normalized with the survival rate of naïve hosts (right; also see Figure S1 for the naïve treatment regime). The experiment was conducted in two replicates for all generations. A cyclical pattern of a drop in survival rate is observed after every two consecutive wasp treatments. After the sixth generation, three consecutive generations show a significant decline in survival. Error bars represent the standard error of the mean. A one-way ANOVA with the Brown-Forsythe test was conducted. **p=0.001, ***p=0.0002.

In parallel, unstressed sibling F_1_ embryos (0-24 hours) were taken and allowed to grow into adult flies to obtain the F_2_ generation. However, some batches of 0-24 hour F_1_ embryos were collected and infected at the second instar larval stage, as described for the experienced treatment group, to determine the percent survival rate of naïve hosts in response to the parasitic stress. An identical method of embryo collection to obtain the subsequent unstressed generation and exposure of some batches to parasitic stress was performed for ten generations based on the scheme presented in Figure S1A, giving rise to the naïve treatment group.

The number of pupae and escapee flies was counted for both naïve and experienced treatment groups in every generation. The number of escapee flies was divided by the number of pupae in the corresponding vial to determine the percent survival rate of the host. The survival of experienced hosts relative to naïve hosts was used to determine the statistical significance and generate bar graphs using GraphPad Prism 7. The number of replicates and details of the statistical test used are indicated in the figure legends.

### Parental contribution

For parental contribution, we exposed second-instar *Drosophila* larvae to parasitoids and carefully separated the resulting virgin females and males that managed to escape. These male and female escapees were referred to as “E_1_” because of their first exposure. Subsequently, we mated the E_1_ males and females with naïve female virgins and males, respectively. Embryos were collected for 24 hours from mated E_1_ male (paternal lineage) and E_1_ female (maternal lineage) escapees to check the parent-specific contribution. Progenies from both the E_1_ lineages were subjected to wasp infection at 24-48 hours. Wasps (10 males and 10 females) were allowed to infect for 24 hours. After infection, larvae were allowed to grow until they emerged as escapees (E_2_) or adult wasps (see schematics in Figures 2B and S2A). The number of escapees with melanized wasp eggs, referred to as melanotic capsules hereafter, was recorded to calculate the survival rate. The melanotic capsule containing male and female escapee flies were used further for setting up crosses to the assess paternal and maternal contribution of the parasitic stress memory to the next generation. This experimental approach was carried out for five consecutive generations (from E_1_ to E_5_), enabling us to assess and compare their relative success when compared to a control group of naïve individuals (Figure S1B) and an experienced group of individuals of bilineal origin (Figure 2A). The experiment was performed in six biological replicates, and a 10:10 male-to-female ratio was maintained throughout.

**Figure 2.**
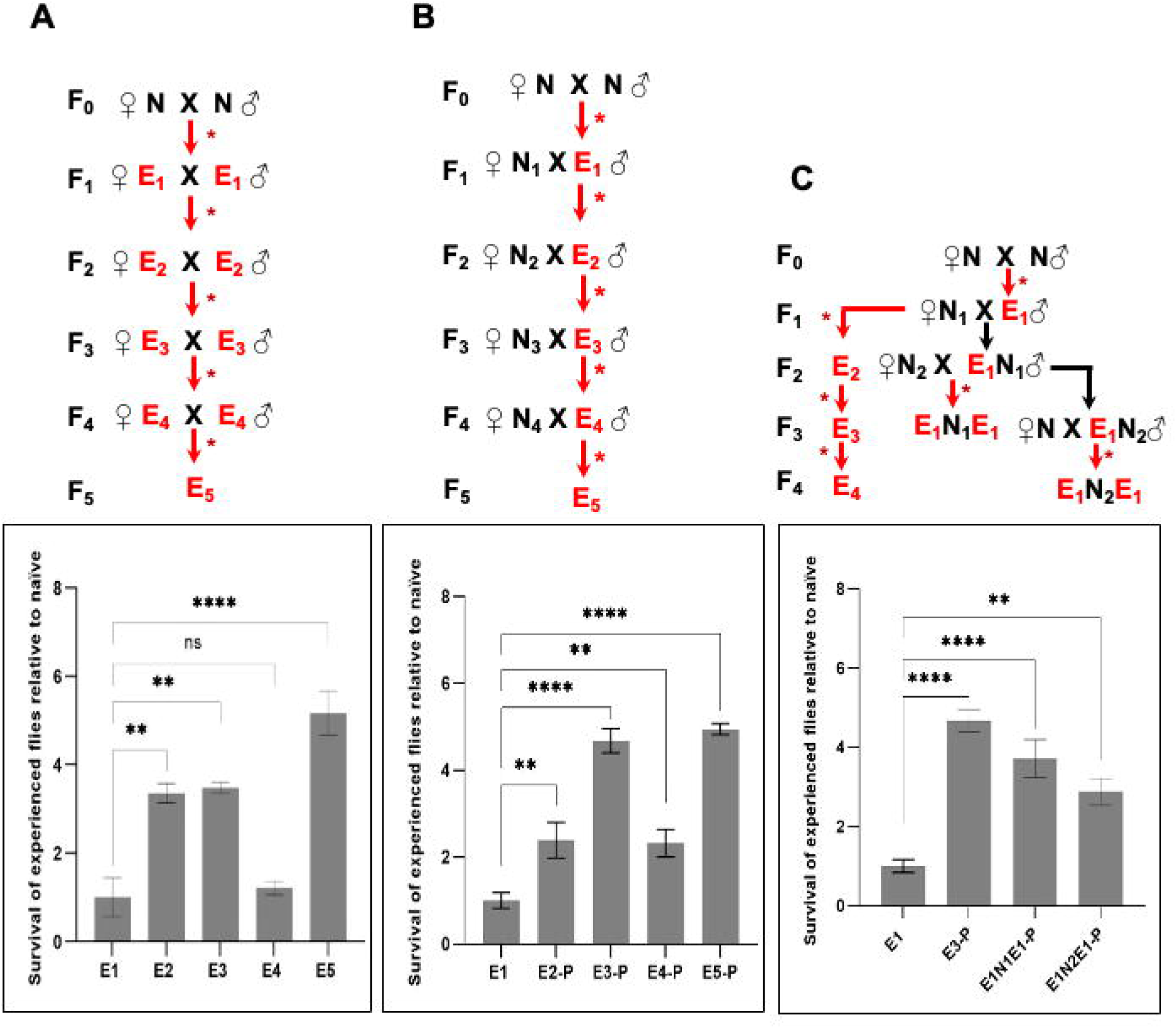
Parental contribution to the parasitic stress memory. **(A)** The F_1_ male and female escapee flies (E_1_) exposed to the parasitic stress at the second instar larval stage were mated. Their progeny were exposed to the parasitic stress to get treatment group E_2_. A similar treatment was repeated every generation for five generations to attain the bilineal inheritance of parasitic memory. The relative survival rate of escapees (E_1_ to E_5_) compared to first-time exposed naïve flies (N_1_ to N_5_) in each generation is plotted in the bar graph. See figure S1B for the treatment regime of naïve flies. The data is from three biological replicates. **(B)** F_1_ males (E_1_) exposed to parasitic stress at the second instar larval stage were mated with naïve virgin female flies. Their progeny were exposed to the parasitic stress to get treatment group E_2_. A similar treatment was repeated every generation for five generations to attain paternal inheritance of parasitic memory. The data is from six biological replicates. **(C)** F_1_ males (E_1_) exposed to parasitic stress at the second instar larval stage were mated with naïve virgin female flies, and their embryos were divided into two groups. One group was exposed to the parasitic stress to obtain a repeatedly exposed legacy (two-time exposed generation, E_2_), and the other group was allowed to grow without any parasitic stress to obtain a one-generation skip legacy (E_1_N_1_). A two-generation skip, for instance, is referred to as E_1_N_2_. This was repeated for four generations to attain paternal inheritance of parasitic memory in a transgenerational manner. The data is from six biological replicates. A schematic of the treatment regime is shown at the top of each bar graph. The red arrows with an asterisk represent parasitic stress, while the black arrows represent the omission of parasitic stress for the corresponding generation. E, experienced escapee; N, naïve. The number in the subscript represents the generation of treatment. For all experiments, we conducted a one-way ANOVA with Brown-Forsythe and Welch’s multiple comparison test. Error bars represent the standard error of the mean. *p=0.0105, **p=0.0088, ***p=0.0001.

### Transgenerational parasitic stress

For the transgenerational regime, once-exposed males and females (E_1_) were collected separately and crossed with naïve virgin females and males, respectively. The F_2_ progeny is referred to as E_1_N_1_ for simplicity. The E_1_N_1_ males (paternal lineage) and E_1_N_1_ females (maternal lineage) were then collected and mated with naïve virgin females and males, respectively (see schematics in Figures 2C and S2B). The F_3_ embryos were collected for 24 hours, and 48 hours AEL were infected by wasps (10 females and 10 males) for 24 hours. Melanotic capsule containing progenies were collected and counted for survival success. This experimental approach was carried out for four generations to assess the transgenerational effect. The control group in all experiments was exposed to wasps only once (Figure S1B). The experiment was performed in six biological replicates, and a 10:10 male-to-female ratio was maintained throughout.

### Immune Induction and immunostaining

Immune induction was done as described previously^14^. In brief, the second instar larvae from naïve parents and exposed parents were poked by a sterile needle at their posterior region in 1X PBS and were then transferred to fresh food vials. After 24 hours, larvae were scooped out of the food vials, washed twice with 1X PBS to remove food remnants, and washed once with 70% ethanol for surface sterilization. The larvae were then transferred to ice-cold 1X PBS until dissection. Haemolymph from a single larva per well was collected and allowed to settle at the bottom of a 4 mm well slide. Cells were then fixed with 4% formaldehyde for 20 minutes, followed by three washes with 1X PBS-T (0.5% Triton-X) for 5 minutes each. After blocking in 1% BSA, cells were stained using the primary antibody anti-myospheroid (DSHB #CF.6G11, 1:500 dilution), a lamellocyte marker, since lamellocytes appear after an immune challenge. After washing three times with 1X PBS-T (0.3% Triton-X), cells were incubated with Alexa Fluor® 647 AffiniPure goat anti-mouse IgG secondary antibody (115-605-003, 1:1000 dilution). Finally, the cells were washed three times with 1X PBS-T (0.3% Triton-X) and stained with DAPI for nuclei. Scanning of each well was done using the Zeiss LSM 880 confocal microscope using the 40X oil immersion lens. For the quantification of lamellocytes, the entire well was scanned, and images were processed by ImageJ Version 1.53c (Fiji). DAPI for total cells and anti-myospheroid for lamellocytes were used to count the cell number in ImageJ.

## RESULTS

### Effect of multigenerational parasitic stress on host survival

To assess the effect of multigenerational parasitic stress on host survival, we designed experiments where fly larvae were repeatedly exposed to wasp for ten generations, resulting in an experienced treatment group with a history of infection (E_1_ to E_10_) (Figure 1B). On the other hand, larvae were newly exposed to wasp infection in every generation for ten generations, giving rise to the naïve treatment group (N_1_ to N_10_) (Figure S1A). We considered all the flies that emerged after infection, with and without the melanotic capsule, to calculate the survival rate of the host. The escapees without the melanotic capsule were taken into consideration for two reasons. First, the fly larvae can escape wasp infection not only by encapsulating the wasp egg (physiological defenses) but also by employing a rolling strategy (behavioral defenses)^39,40^. Second, since it has been shown that in *D. melanogaster* the melanization rates are low^48^, which can be attributed to the low haemocyte load of *D. melanogaster* as compared to other *Drosophila* species^49^, the absence of a melanotic capsule doesn’t necessarily indicate a lack of infection. However, only melanotic capsule containing escapee flies were taken as parents to obtain the subsequent generations to ascertain infection.

We observed that infecting the progenies of F_1_ escapee flies (E_1_) at the larval stage i.e., E_2_, resulted in a significant increase in survival after infection (Figure 1B) when compared to the survival after infection of progenies of F_1_ naïve flies (N_1_) at the larval stage (Figure S1A). A similar increase in survival was also observed when the progenies of F_2_, F_4_, F_5_, and F_9_ escapee flies (E_2_, E_4_, E_5_, and E_9_) were infected as compared to N_2_, N_4_, N_5_ and N_9_, respectively, except for the progenies of F_3_ (E_3_) generation. Overall, we see a cyclical increase and decrease in the total number of successful escapees. These results indicate that the progenies of the parents exposed to the parasitic stress acquire better survival capability for the subsequent attacks.

### Male parents effectively pass the parasitic stress memory to their progeny

We investigated whether both parents equally contribute survival advantage to the progeny by allowing either the male or female escapee flies to give rise to the subsequent generation and thereby contribute to the parasitic stress memory. Exposed male flies (E_1_) were mated with naïve female flies to examine the paternal inheritance of parasitic stress memory (Figure 2B). On the other hand, the exposed female flies (E_1_) were mated with naïve male flies to examine maternal inheritance of the parasitic stress memory (Figure S2A). The experiment was carried out for five generations. As a control, naïve parents (Figure S1B) and experienced parents of bilineal origin (Figure 2A) were taken. Interestingly, male parents were able to inherit the survival advantage to their progeny in every generation when the parasitic stress was given repeatedly, whereas the female parents could not inherit the memory beyond one generation (Figures 2B and S2A). While progenies from the experienced mother showed better survival in only one generation compared to the once-exposed control, the survival advantage was more than twofold in the progenies of experienced fathers in the subsequent generations upon repeated exposure.

We further examined if increased survival is a result of increase in egg lay, such as when a stressed adult female fly tends to lay more eggs once the stress is removed. We checked the fecundity of progenies from stressed parents and found no significant difference in the fecundity of the progenies from stressed parents compared to the naïve flies (Figure S3).These results indicate differences in the perception of and response to the parasitic stress of male and female flies.

### Parasitic stress memory is transgenerational

We further explored whether the parasitic stress memory is transgenerationally inherited. We set two groups of experiments where in one group, E_1_ males were mated with naïve virgin females to obtain the paternal lineage, and in the other group, E_1_ virgin females were mated with naïve males to obtain the maternal lineage. Progenies from both lineages were collected without any wasp exposure to get unexposed male or female (E_1_N_1_) progenies. E_1_N_1_ males from paternal lineage and E_1_N_1_ females from maternal lineage were then mated with naïve females and males, respectively, and their progenies were exposed to wasps at the second instar larval stage. These larvae, named E_1_N_1_E_1_ (grandchildren of once-exposed males or females), were allowed to grow in standard conditions. Unlike female grandparents, male grandparents successfully inherited the parasitic stress memory, which is manifested as the survival advantage, to their grandchildren as compared to the control (once exposed) (Figures 2C, S1B, and S2B). Moreover, we observed that the parasitic stress memory was inherited beyond two generations (E_1_N_2_E_1_). These results suggest transgenerational inheritance of the parasitic stress memory to subsequent generations via the male germline.

### Adaptive memory is passed on as cellular immunity

*Drosophila* exhibits a cellular immune response upon wasp attack. It possesses three types of haemocytes engaged in the immune response: plasmatocytes, which constitute 95% of the total haemocytes and eliminate pathogens and injured cells; crystal cells (5%); and lamellocytes, which are rarely observed in healthy larvae. Lamellocytes emerge following plasmatocyte differentiation in response to any foreign immune challenge and are deployed to encapsulate the pathogen, depriving it of oxygen and nutrients^41,44,50,51^. Therefore, we speculated that the survival advantage observed in the progenies of stressed parents is due to an enhanced cellular immune response. In order to test that, we induced the progeny of exposed parents using a sterile needle to mimic the wasp attack and measured the total number of haemocytes and lamellocytes 24 hours post-induction. Progenies of exposed males showed an elevated number of lamellocytes compared to the control (induced larvae from unexposed parents) in all four generations (Figure 3). One-time wasp exposure results in an almost four fold (18% of the total hemocytes) increase in the lamellocyte percentage compared to the naïve-induced larvae (4%). Similarly, the larvae obtained from parents stressed for two and three generations showed a significant increment in the lamellocyte percentage. While the progeny of the exposed females show a slight increase in the lamellocyte number compared to the control, it is not equivalent to the progeny obtained from exposed male parents (Figure S4). These results indicate that the survival advantage observed after multiple generations of parasitic stress is correlated with the enhanced cellular immune response of the hosts.

**Figure 3.**
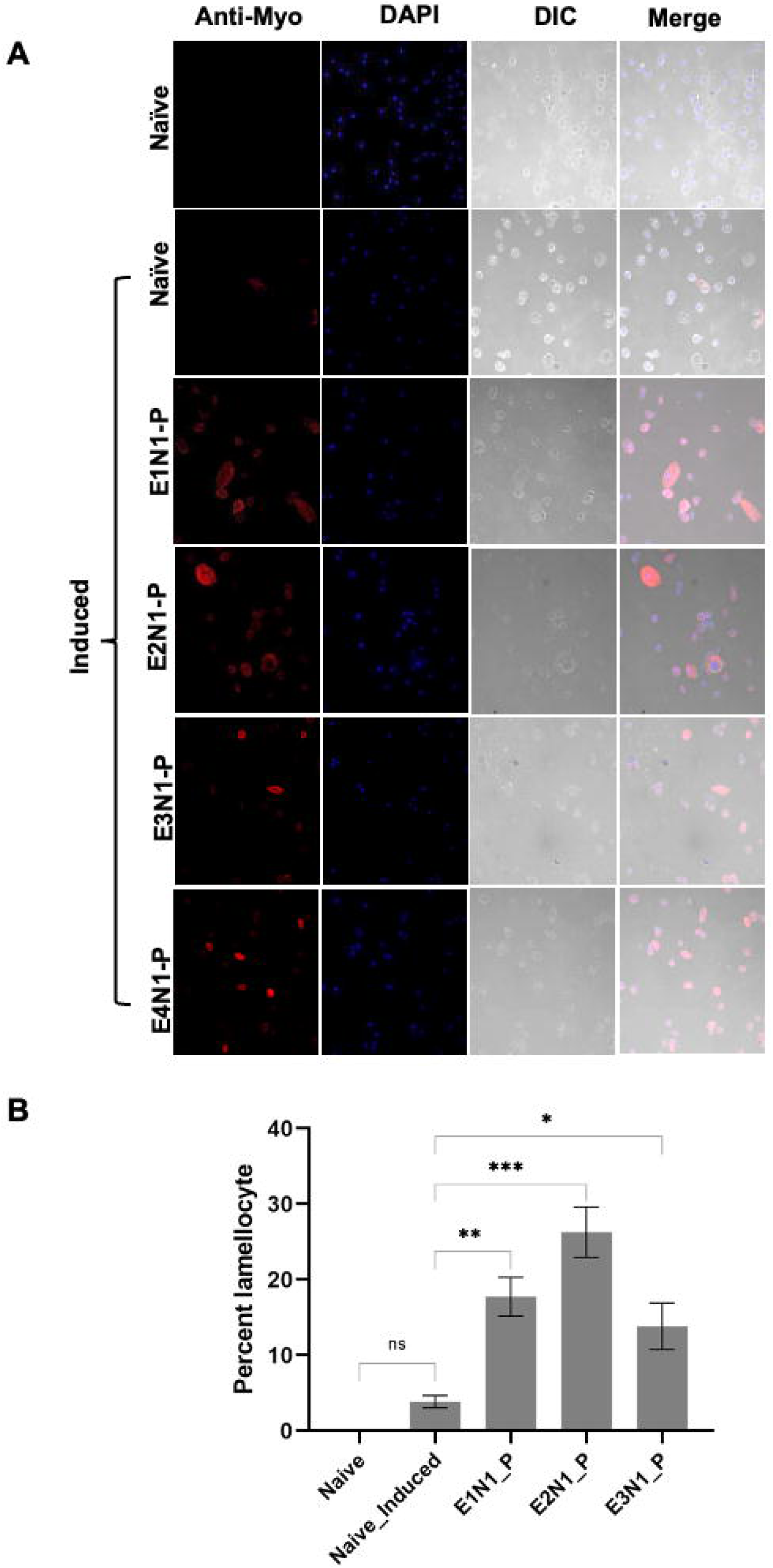
Cellular immune response to parasitic stress in progenies of male parents. **(A**) The panels represent images of circulatory hemolymph in the third instar larvae of experienced male parents. In the first panel, third-instar larvae from naïve parents were bled out and stained with anti-myospheroid (red) as a lamellocyte marker and DAPI (blue) to stain nuclei. The second panel displays hemocytes from third instar larvae that were mechanically induced at the second instar larval stage to mimic a wasp attack. In the third panel, hemocytes from third instar induced larvae of one-time experienced male parents (E_1_N_1_-P = larvae from male parents exposed to wasps once) are shown. The myospheroid-positive cells in this panel are either fully developed lamellocytes (big and elongated) or cells committed to developing into lamellocytes. The presence of lamellocytes in the hemolymph indicates an elevated cellular immune response. The fourth panel, E_2_N_1_-P, represents hemocytes from induced larvae that come from two consecutive generations exposed to wasps through the male parent. The fifth panel, E_3_N_1_-P, displays hemocytes from induced larvae that come from three consecutive exposed generations, and the sixth panel, E_4_N_1_-P, shows hemocytes from induced larvae that come from four consecutive exposed generations. **(B)** Quantification of Lamellocytes in Larval Hemolymph across Generations from the Paternal Lineage. We quantified the number of lamellocytes in larval hemolymph at all generations from the paternal lineage. A slight change in induced naïve larvae is observed compared to naïve larvae. One generation of exposure shows an almost four-fold increase compared to induced naïve larvae. Two generations (E_2__P) and three generations exposed (E_3__P) progeny also show almost five- and three-fold increases in myospheroid-positive cells, respectively. The experiment was conducted in three replicates, and a one-way ANOVA with Brown-Forsythe and Welch’s multiple comparison test was performed. Error bars represent standard errors, with *p=0.0105, **p=0.0088, ***p=0.0001.

## DISCUSSION

Organisms are constantly engaged in the evolutionary arms race. Success in multi-organism interactions, such as host-parasite interactions, depends on how strong or prepared the defense system of the host is and how sneaky the parasite is to escape the host defense arsenals. Innate immunity in insects is one major deterrent for parasites and pathogens during the embryonic and larval development stages. If the progeny is alerted by the parental message in the form of epigenetic memory, it may be a crucial factor to the host’s advantage. Immune priming is defined as a phenomenon wherein the parental experience of infection results in resistance to infection in the offspring^4^. In this study, we demonstrate that in response to continuous parasitic stress by wasps, the fruit flies produce progenies that can withstand the stress better, perhaps by developing a better defensive system, either physiological or behavioural.

It has been shown previously that cohabiting fruit flies with adult female wasps results in intergenerational immune priming of fly offspring; that is, the immediate offspring or larvae of cohabitated flies show increased survival after wasp infection^14^. The study also showed an increased survival rate correlated with enhanced production of lamellocytes, a type of immune cell in flies. Consistent with this, we also see immune priming of the offspring of escapee flies that survived the wasp attack, although our study represents a case of immune priming due to a direct infection and not cohabitation. However, further investigation would be needed to decipher if the immune priming of the offspring observed in this study after parental infection is transmitted via the epigenetic route of inheritance.

We further asked if both parents contribute to parasitic stress memory. Our results show that only the male parent can transmit the memory to subsequent generations, inter- and transgenerationally. Female parents do not show successful transmission of the survival advantage upon repeated exposure to the wasp. This indicates that the cumulative memory of wasp exposure is either detrimental to female germline development, as previously shown^34^, or that the memory is not maintained during female germline development. Our study shows that after two subsequent exposures in the maternally inherited lineage, the third generation does not maintain the memory. This indicates that the female parents do not choose memory maintenance when the exposure is continuous. Perhaps they can protect their progeny through behavioural strategies. Female flies exhibit behavioural defenses, such as egg lay avoidance in the presence of female wasps, alcoholic food preference for egg lay, and change in mating behaviour^30–32,52^. While male parents have less chance to provide direct defense to their progeny, the only way to convey their experience is via the germline. It is also plausible that the two parents might have evolved unique techniques to articulate the memory of their negative experience with a parasitic attack.

Increased lamellocyte numbers show that the host-parasitoid interaction induces transgenerational immune priming, which helps the progeny be ready for the upcoming wasp attacks. However, in our experiment, the fourth generation shows a decrease in both survival and lamellocyte numbers (Figures 2B and 3B). Although it remains elusive without further experiments, we speculate that the cellular defense has its own cost, and the cumulative memory of three generations makes the host weak, leading to increased lethality. Nevertheless, the memory of their experience can be inherited via epigenetic changes in the germline, which calls for further exploration.

In conclusion, we show that parasitic stress memory is transgenerationally inherited through the male germline in *D. melanogaster*. This draws attention towards future studies on the possibility of multigenerational inheritance of past experiences of biotic stress via germline-mediated epigenetic mechanisms. How such a memory is transmitted through the sperm remains to be explored. While such studies will show how the parental history of wasp infection at the pre-adult life stage of flies imparts a survival advantage to the subsequent generation without any social communication, what also remains to be explored is how widespread such epigenetic inheritance mechanisms are across the animals and how many kinds of stress are covered by them. Finally, it would be of interest to know if different mechanisms exist for different stresses or if these are broad-natured defense mechanisms for a variety of anticipatory harms.

## ACKNOWLEDGMENTS

We thank Indira Paddibhatla for introducing the *Drosophila–Leptopilina* system to our lab. We thank Sharath Chandra Thota and Gottivedu Jyothirmai for helping in experiments. We thank N.R. Chakravarthi, C. Subbalakshmi and B. Suman for technical assistance in imaging facility. We thank the staff of the *Drosophila* Laboratory at the Centre for Cellular and Molecular Biology for providing technical assistance. S.K. thanks the Department of Science and Technology (DST) for the INSPIRE fellowship. R.S thanks Council of Scientific and Industrial Research (CSIR) for fellowship. R.K.M. laboratory is supported by CSIR and SERB-DST.

## AUTHOR CONTRIBUTIONS

S.K., R.S., and R.H. executed the study and analysed the data. S.K. did the multigenerational parasitic stress experiment in both parents, R.S. and R.H., did paternal and maternal intergenerational parasitic stress experiment, R.S. did maternal and paternal transgenerational parasitic stress, immune induction and imaging. S.K., R.S., R.H. and R.K.M. wrote and edited the manuscript. R.K.M. conceived the project and supervised the study. All authors read and approved the final manuscript.

