## supplemental figures for "Multigenerational inheritance of parasitic stress memory in *Drosophila melanogaster*"

### Figure S1

A

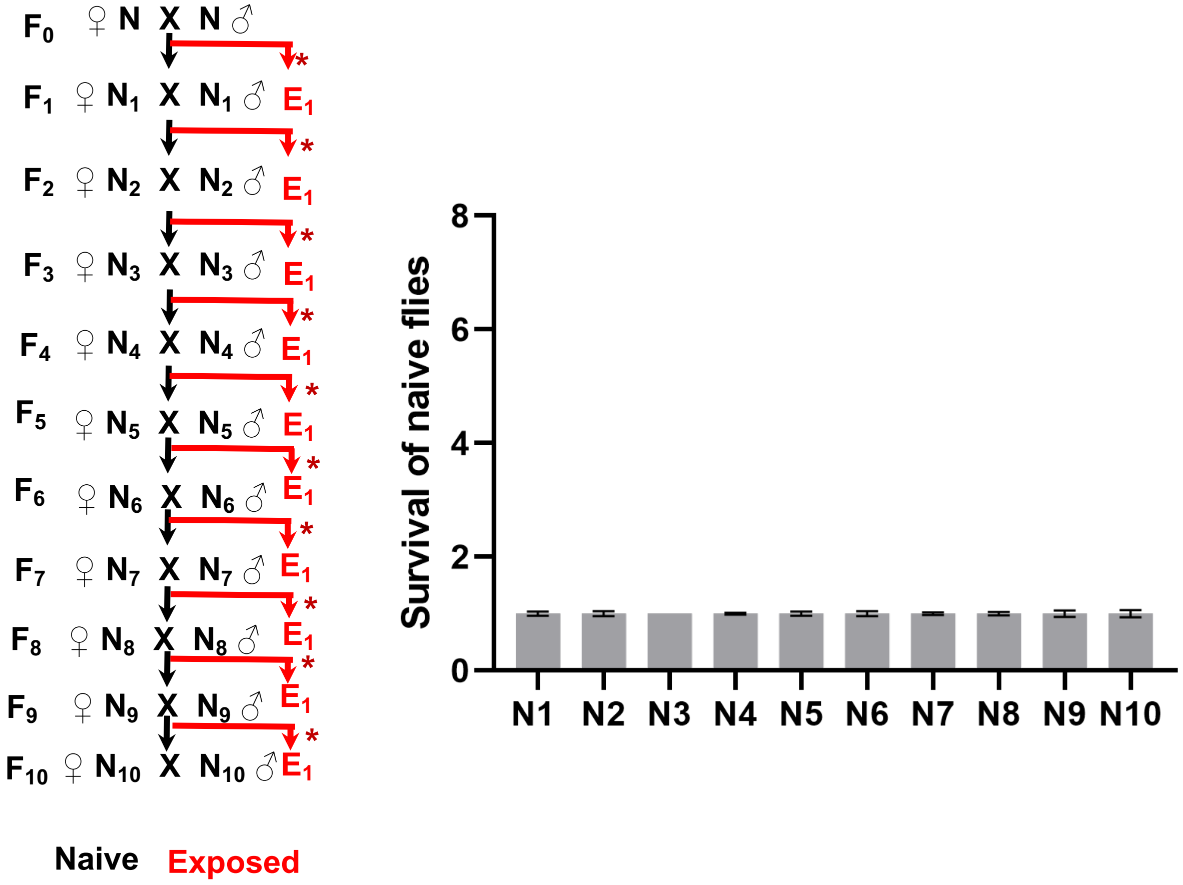


**Figure S1A:** Parasitic stress regime for ten generations (left) and the corresponding survival rate (right) of the flies in every generation. Naïve flies were propagated parallel to all the multigenerational experiments, intergenerational and transgenerational, and used as once-exposed control in every generation. Normalized value of survival rate of naïve flies up to ten generation is shown in the graph.

### Figure S1

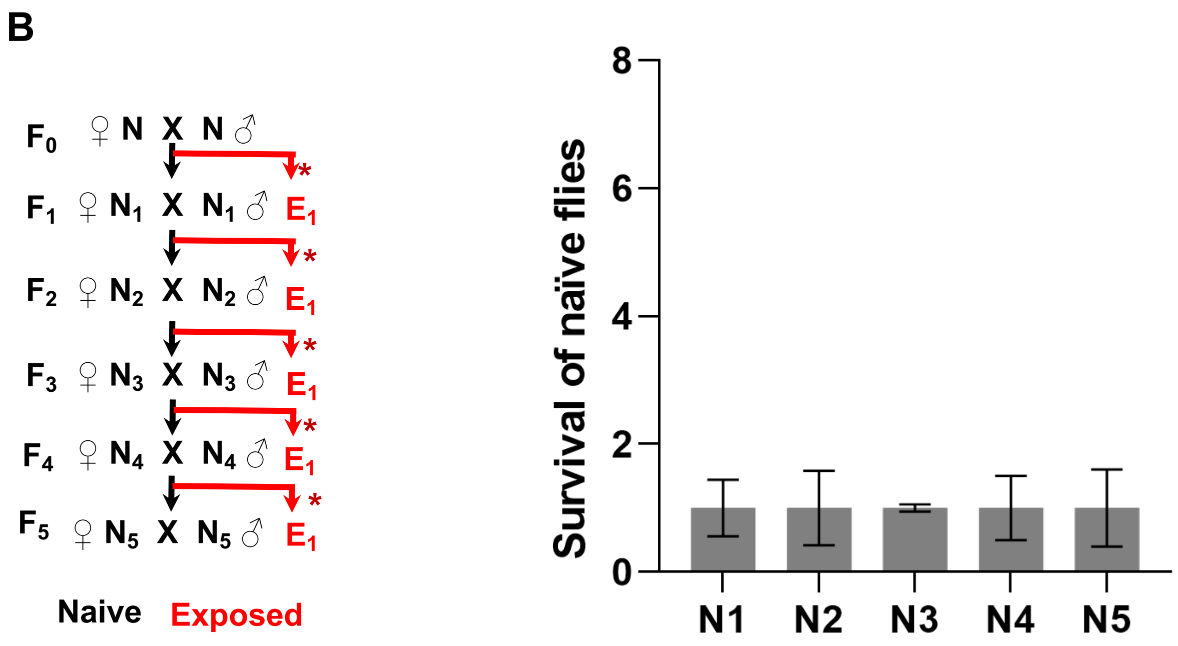


**Figure S1B:** Parasitic stress regime for five generations (left) and the corresponding survival rate (right) of the flies in every generation. Naïve flies were propagated in parallel to all the multigenerational experiments, intergenerational and transgenerational, and used as once-exposed control in every generation. Normalized value of survival rate of naïve flies upto five generations is shown in the graph.

### Figure S2

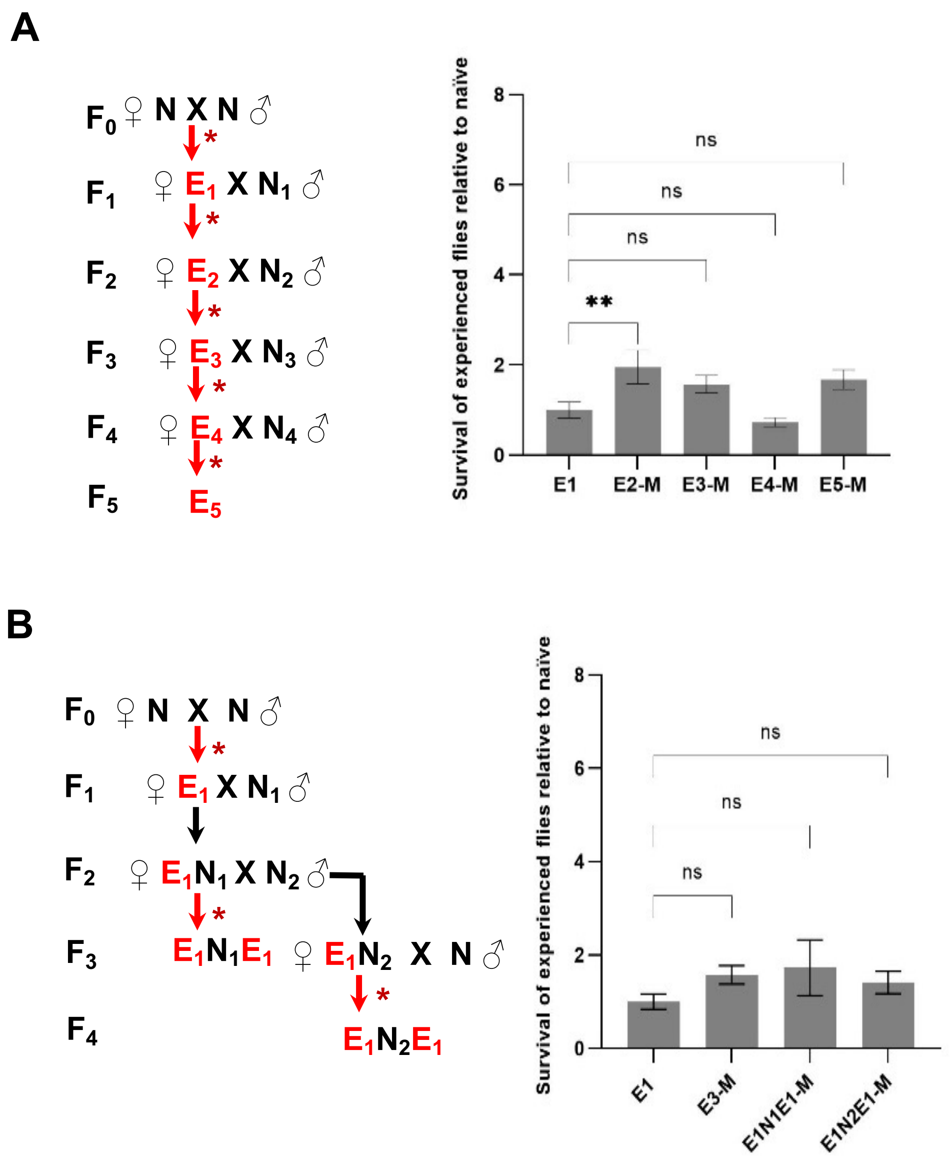


**Figure S2. (A)** F1 females ($E_1$) exposed to parasitic stress at the second instar larval stage were mated to naïve male flies. Their progenies were exposed to the parasitic stress to get treatment group $E_2$. A similar treatment was repeated every generation for five generations to attain maternal inheritance of parasitic memory. **(B)** $F_1$ females ($E_1$) exposed to wasps at second instar larval stage were mated to naive male flies. Their embryos were divided into two groups. One group was exposed to the parasitic stress to get the repeatedly exposed legacy (two-time exposed generation, $E_2$) and the other group was allowed to grow without parasitic stress to get one-generation skip legacy ( $E_1N_1$ ). A two-generation skip is referred to as $E_1N_2$. This was repeated for four generations to attain maternal inheritance of parasitic memory in a transgenerational manner.

The treatment regime is shown on the left side of the bar graph. The red arrows with an asterisk represent parasitic stress, while the black arrow represent the omission of parasitic stress for the corresponding generation. E, experienced escapee; N, naïve. The number in the subscript represents the generation of treatment. The data is from six biological replicates. A One-way ANNOVA with Dunnet's multiple comparison was done for all experiments. Error bars represent the standard error of the mean. *p=0.0105, **p=0.0088, ***p=0.0001.

### Figure S3

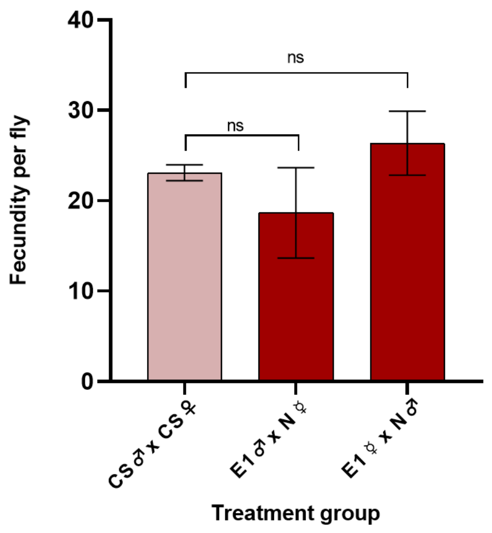


**Figure S3:** Fecundity per fly per day. Eggs from each category – control (CS males X CS females), experienced females mated with naïve males (♀ $E_1$ X $N_1$ ♂ ), and experienced males mated with naïve females (♀ $N_1$ X $E_1$ ♂) – were counted for five days. Total number of eggs were divided by the number of days and the value is plotted on graph for each category. There is no significant change in any category compared to control (♀CS X CS ♂).

### Figure S4

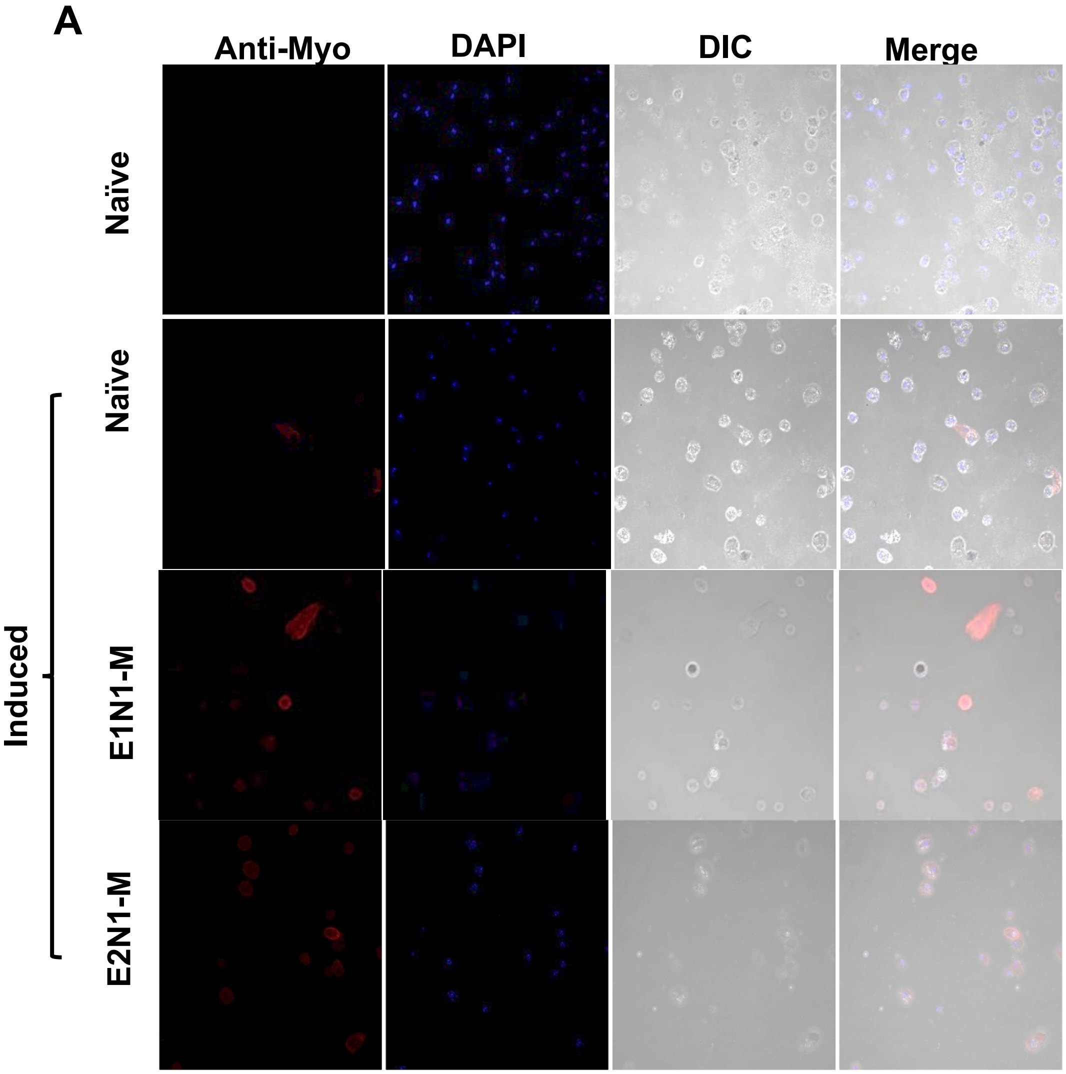

### Figure S4

**B**

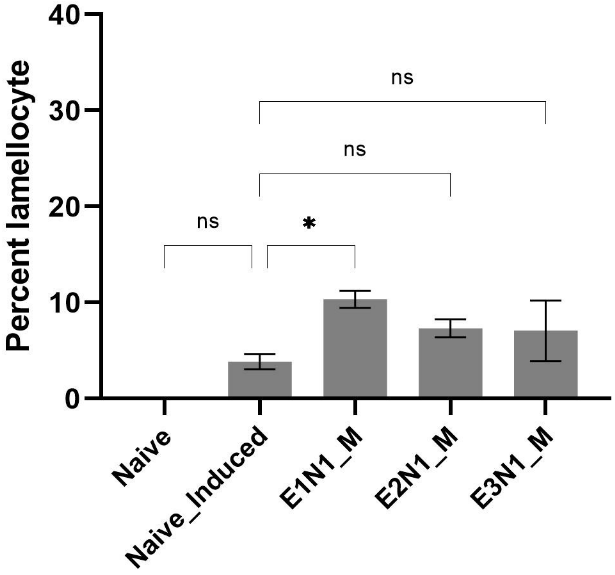


**Figure S4. Cellular immune response to parasitic stress in progenies of experienced females. (A)** Representative images of circulatory hemolymph of third instar larva of experienced female parent. $E_1N_1$_M = female parent was wasp exposed, M - maternal, $E_2N_1$_M = two consecutive generations from wasp exposed female parent. Red-myospheroid positive lamellocyte, blue =DAPI. (**B)** Quantification of lamellocyte in larval hemolymph at all generations from maternal lineage. One generation exposure shows almost two-fold increase compared to induced naïve. Two generations ($E_2N_1$) and three generations ($E_3N_1$) exposed does show a slight increase, but not significant change compared to naïve induced.
